# Effect of level of autistic traits on motion inspection time performance in centre-surround suppression tasks

**DOI:** 10.1101/2022.06.06.495054

**Authors:** Andrew Gubko, David Crewther

## Abstract

The mechanisms underlying visual anomalies observed in individuals with Autism Spectrum Disorder (ASD), presenting in the form of impaired global visual perception and enhanced local perception have long been debated. Researchers have sought to investigate the phenomenon of surround suppression, the inhibitory mechanism responsible for filtering visual noise in high contrast stimuli of increasing size. Mixed results have shown impaired gain control reduces surround suppression allowing for improved motion inspection in certain conditions, along with the possibility that receptive field size and population varies among the ASD population producing size-specific alteration in motion inspection. The present study aims to investigate whether the level of autistic traits predicts motion inspection time performance in a random dot kinetogram (RDK), a function of stimulus size and contrast. A total of 12 typically developing (TD) participants were screened and grouped by AQ score to indicate level of autistic traits (low, medium and high). Results revealed that the high AQ group’s motion inspection time significantly outperformed the mid AQ group for small and large central RDK of low contrast. This result suggests an interaction between temporal and spatial processing may have masked the effects of gain control and impaired surround suppression in higher contrast conditions, often seen in previous studies.

## INTRODUCTION

Autism spectrum disorder’s (ASD) atypical sensory processing originates from Kanner’s (1943) early accounts regarding children with delayed speech, fixation for spinning objects and abnormally specific memory. In an attempt to understand the atypical social and sensory abnormalities present among the ASD population researchers have promptly investigated the various facets of their neurological mechanisms. Anomalies and alteration in sensory development has shown to have a high prevalence with an estimate of 69–95% in children with ASD (Manelis-Baram et al., 2021).

Self-assessed questionnaires are often utilised in studies to explore varying levels of performance as a function of autistic traits that an individual displays (Wykes, Hugrass & Crewther, 2018; Jackson et al., 2013). Robertson and Simmons (2012), found that both hyper sensitive and hypo-sensitive responses were evident in individuals with higher levels of autistic traits (established with the use of the Autism spectrum quotient (AQ of Baron Cohen et al., 2001). Sensory studies investigating ASD have often provided findings for clinical populations, however, without screening control groups it is not suitable to normalise these results without recording or grouping individuals by level of autistic traits. High vs low level of autistic traits in typical developing populations has allowed for insight into how surround suppression, abnormal magnocellular functioning and binocular rivalry occur across the autism spectrum rivalry (Jackson et al. 2013; Robertson & Simmons, 2012; Wykes, Hugrass & Crewther, 2018).

Visual processing is facilitated through two cortical visual pathways: the dorsal stream branching through the parietal lobe and the ventral stream projecting to the temporal lobe (Spencer et al. 2000). The dorsal stream processes spatial awareness and locates how and where stimuli move in the surrounding environment, as well as informing visually driven actions. The ventral stream allows for processing of the form and identification of visual stimuli (Kravitz et al., 2013). The argument that these are not independent processes emphasises the complexity in identifying how anomalies in visual processing occur in a spectral disorder such as ASD (Kravitz et al., 2013; Milner, 2017). Lesion studies have demonstrated that impairment in the ventral pathway promotes compensatory mechanisms in the dorsal pathway (Bridge et al., 2008).

Current understanding of visual pathways focus on how information is integrated rather than solely on their respective inputs and outputs, thus the dorsal pathway contribution in visuomotor control currently labels it as the “how” pathway rather than its former label and function of the “where” pathway (Freud, Plaut & Behrmann., 2016; Rauschecker 2018). The medial temporal (MT) is a crucial component of the dorsal pathway, as it relays spatial information to the parietal and temporal lobes for further processing of visually guided movements and visual memory (Dutton 2003). The MT receives inputs from the lateral geniculate nucleus (LGN) and provides information about movement and position of objects relative to the space it occupies. The MT is highly innervated by magnocellular input from the LGN. Magnocellular neurons are rapidly adapting large cells that are sensitive to motion and depth in contrast whilst parvocellular neurons are slowly adapting and sensitive to colour and fine details.

Observing behavioural recounts of ASD individuals displays how aspects of the dorsal pathway operate differently compared to typically developed individuals (Kanner, 1943). The difficulties observed in global visual perception and facial recognition in ASD can somewhat be compared to an underdeveloped visual system seen in early childhood. Studies mention within the first few months of life, aspects of the retinopulvinar-MT pathway are abundant but under irregular conditions are unable to prune these connections and mature the LGN pathway to dominate the visual cortex (Bridge, Leopold & Bourne 2016; Mercier et al. 2016). A retinopulivinar-MT pathway that persists through adulthood may be the underlying reason for vulnerability noticed in the dorsal pathway in ASD, as ganglion cells entering the Plm terminate on neurons that project to the MT (Braddick, Atkinson & Wattam-Bell, 2003; Pellicano & Gibson, 2008; Bridge, Leopold & Bourne, 2016). This prevents mature development of V1 to MT, influencing integration of global perception. Visual literature mentions that local motion signals produced by V1 are spatially integrated into MT neurons which comprise of larger receptive fields, required for global motion perception (Furlan & Smith, 2016; Van der Hallen et al., 2019). Sensitivity to direction of motion within a small region arises from the small receptive fields that occupy the V1. Rather in ASD there is a notable reliance on parvocellular pathway and identifying local over global motion (Boxtel & Lu, 2013).

Tadin et al. (2003) explains that retinal ganglion cells within the visual cortex contain receptive fields that are direction selective due to their centre-surround properties. Their findings suggest that as contrast increases for large stimuli, surround suppression propagates, weakening the ability to detect the motion of central stimuli in conditions with surrounding movement. For large stimuli with low contrast, visibility of central moving stimuli improves due to decreased surround suppression.

The prevalence of centre-surround suppression in motion inspect time performance for individuals with ASD has been studied extensively (Bakroon & Lakshminarayanan, 2018; Shuffrey et al. 2018; Foss-Feig et al., 2013; Schauder et al., 2017; Schwarzkopf et al., 2014). Branching from the research of Tadin et al. (2003) it has been proposed that the size and population of these centre surround receptive fields differ in size or population among a clinical ASD population and manifest perception in the form of delayed or enhanced motion inspection time in surround suppression tasks (Schauder et al., 2017; Schwarzkopf et al., 2014; Foss-Feig et al., 2013; Spiteri & Crewther, 2021). Change in size of receptive fields, population and even firing rate may be due to vulnerabilities in the dorsal pathway discussed previously to compensate for integration of local information in a global manner as proposed by Bridge, Leopold & Bourne (2016).

Studies measuring performance in tasks involving surround suppression, typically use motion inspection time PEST (parameter estimation by sequential testing) to alter difficulty (Tadin et al., 2003), however coherence levels PEST have also been employed (Milne et al. 2002). Tadin et al., (2003) hypothesised that increasing stimulus size in a high contrast condition with short temporal duration would elicit surround suppression in a typically developed population requiring more time to detect global motion. However, the study design if Tadin et al., (2003) study design did not account for surround peripheral conditions. Furthermore parasol cells, providing inputs to the motion pathway, are relatively more abundant in the peripheral retina than the fovea compared with the midget cells which feed into the parvocellular pathway. It has been hypothesised that the increase in parasol cells demonstrates intact motion detection when the motion signals are not embedded in visual noise (Thompson et al. 2007). This is why studies have commonly shown increased surround suppression and weakened orientation perception of moving stimuli in the periphery (Xing & Heeger, 2000). To accommodate for this, our study will also implement a surround peripheral condition to reveal whether surround suppression is reduced peripherally in higher traits of ASD compared with foveal vision.

Past research has presented with largely inconsistent results despite the use of similar study paradigms. Gain control, an inhibitory mechanism present among neurons across the visual system. This mechanism is vital in processing motion of stimuli filtering out irrelevant contrast variations and allows visibility for perception. Some of the results for motion inspection time indicate an enhanced motion perception due to impaired gain control allowing for weakened surround suppression in high contrast conditions (Foss-Feig et al., 2013; Lu & Sperling, 1996). However other researchers oppose this finding suggesting irrespective of contrast, ASD individuals show delayed motion inspection times for small sized stimuli. This demonstrates that larger receptor fields reside within the ASD visual pathway making it more difficult for receptive fields to differentiate smaller coherent dots from one another, whilst medium and large stimuli are more distinguishable (Schauder et al., 2017; Schwarzkopf et al., 2014). These findings are based upon the use of a drifting grating stimuli, a 2 alternative forced (2AFC) task assessing ability to detect whether the stimuli is moving left or right. However, it is discussed that the use of a random dot kinetogram (RDK) task is expected to measure the extent of motion inspection time delay in integrating global processing of local details (Bakroon & Lakshminarayanan, 2018).

The present study aimed to utilise the parameters set for contrast and size used in previous drifting grating research to be used in 4 AFC RDKs to measure whether individuals with higher autistic traits demonstrate longer motion inspection times in small sized stimuli and improved motion inspection time at high contrast irrespective of stimuli size. Motion inspection time Bayesian thresholds were employed along with surround peripheral conditions. The new experimental stimuli draws upon the 2 AFC drifting gratings and gabor patches from previous work in terms of stimulus size and contrast, but instead use inner and outer RDK circles (borderless) with 4 AFC choice of motion direction for the surround or central motion. The aim was to demonstrate whether an individual with high autistic tendency displays intact surround suppression under these conditions.

## METHODS AND MATERIALS

### Participants

Debriefing was conducted along with written informed consent was obtained from all participants who partook in the study. The study was advertised on social media sites (Facebook and Reddit) as well as direct email. Participants who completed the online qualtrics AQ questionnaire were emailed a file along with instructions on how to install Psychopy, perform the task and email results back to the researcher. The AQ scores derived from a 160 question questionnaire grouped the participants into low (>11), medium (12-22) and high (23+) austic trait groups for analysis with motion inspection time scores. Vividness of visual imagery questionnaire (VVIQ) consisting of 6 questions was Selection criteria included: no history of neurological conditions, normal or corrected to normal vision and education/job descriptions were recorded in the questionnaire.

Sixteen typically developed adults (9 male, 3 female) with AQ scores ranging between 13 – 37 (M= 23 years, SD= 3.38 years) participated in the study. One of the 3 high AQ scorers included a participant with diagnosed Autism, and no low AQ was recorded among the sample. In total 4 participants were excluded, two reaching ceiling effect (exceeding 0.5 seconds motion inspection time) due to misunderstanding of the task, one did not agree to complete the questionnaire and one had tested the trials repeated amount of times during the programing phase leading to practice effect.

### EXPERIMENTAL STIMULI AND STUDY DESIGN

The motion tasks involved the use of 4 alternative forced choice (4AFC) random dot kinetogram with 6 conditions. 4 AFC requires identification of motion with 4 choices (up, down, left or right) the 25% chance probability of guessing a trial correctly compared to the former 2 AFC (50% chance) meaning motion inspection time threshold will be achieved quicker. The present study investigated the variables of size and contrast with the use of small/large stimuli in low/high contrast conditions. Similar study design was used based on the work of previous motion tasks that used 2AFC drifting gratings (Tadin et al. 2003; Foss-Feig et al. 2013;Schauder et al. 2017;Shuffrey et al. 2018). Stimuli was created on Psychopy 3 (Peirce et al. 2019).

The six conditions consisted of 30 trials each (180 trials total). A tutorial practice run of 8 trials was conducted with varying size and contrast, upon completing the actual task commenced. The first two trials (large condition) had an outer circle with a radius of 15 degrees and borderless inner with a radius of 6 degrees (see **Figure 1**). The second two trials (small condition) had an outer circle with a radius of 15 degrees and an inner circle of 2 degrees. Both the large and small condition had a low and high contrast version, high being white dots on black background and low contrast being white dots on a grey background. The final condition was a surrounding peripheral condition that had an outer circle of 15 degrees and inner circle of 6 degrees for both low and high contrast.

**Figure 1.**
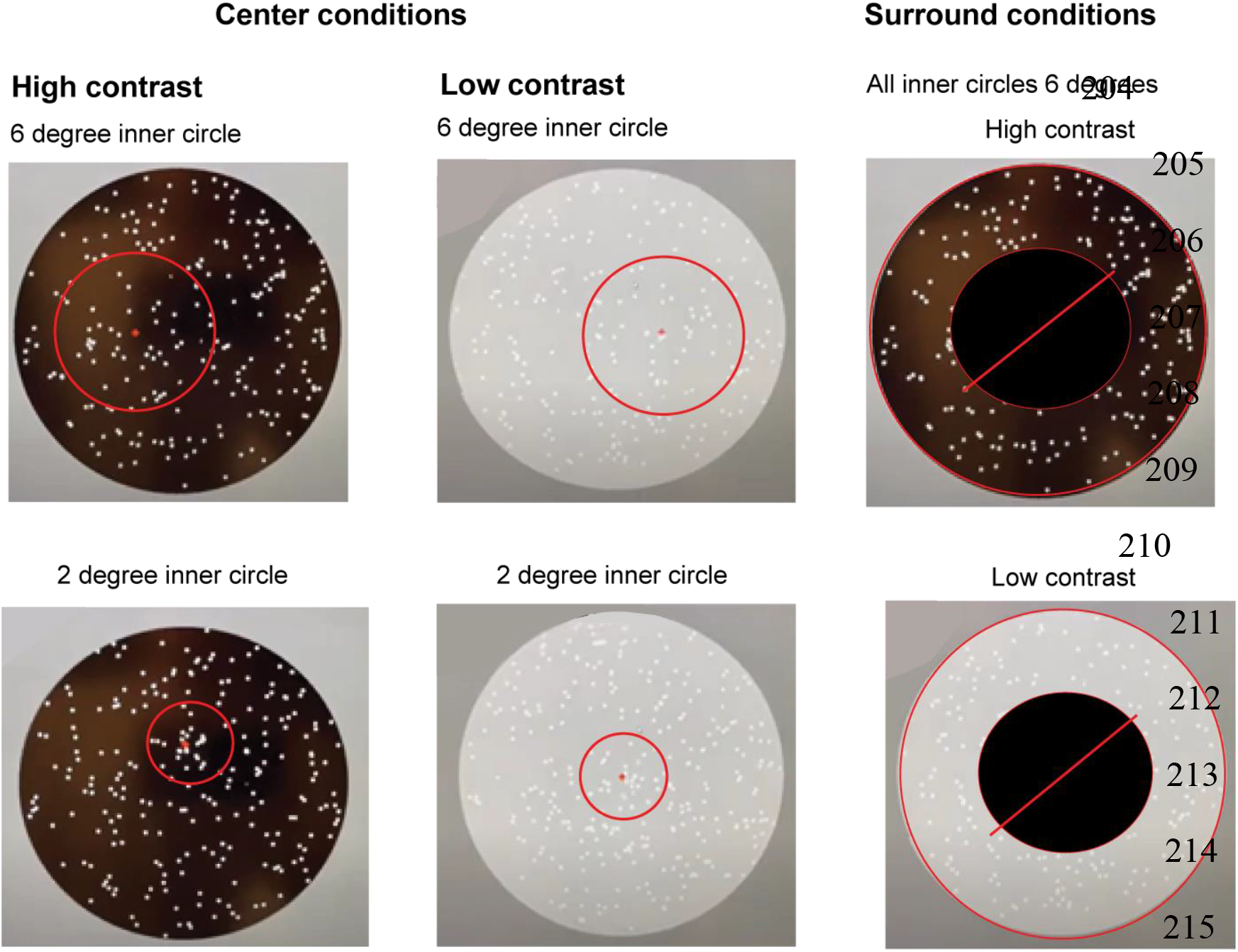
Experiment stimuli conditions. Red circle did not appear during inspection of dot movement it appear prior to dots appearing. Surround conditions do not present with black circle, this is only to demonstrate 6 degree area the participants must ignore. During trial the inner circle in all conditions switched between 6 random locations to ensure spatial uncertainty.

**Figure 2.**
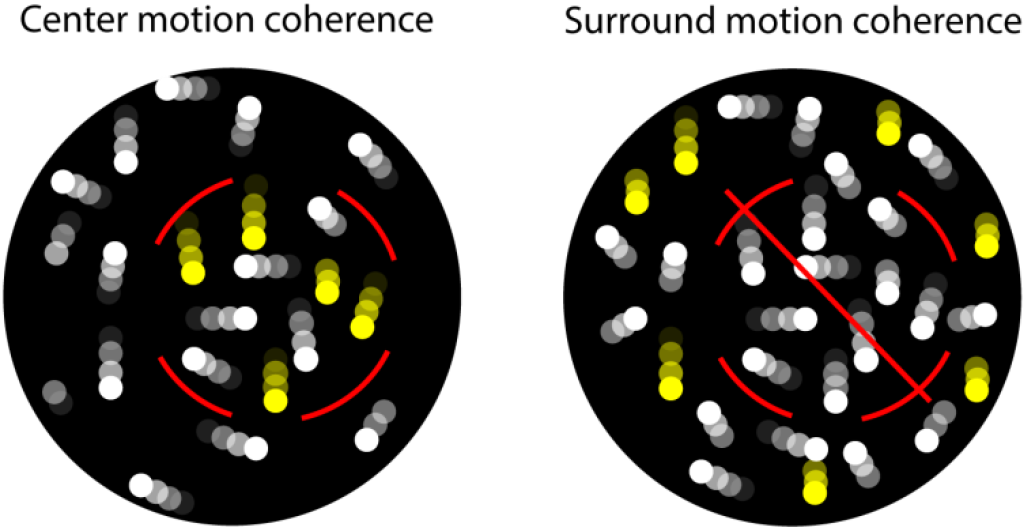
Center and surround motion coherence conditions. Whilst dots were moving with the same constant speed in both regions, the coherence was 50% coherence in the target region (center or surround) while the motion coherence of the dots or outside of region of interest (for both center and surround conditions) were set at 0% to ensure there are no cases where direction of inner and outer dots move together in unison.

The study utilised a QUEST (Bayesian adaptive psychometric method) interleaved staircases design with the stimuli first presenting for 0.5 seconds. Correct identification of the direction the inner circle dots are moving, will mean the time the stimuli is presented will drop for the next trial, an incorrect response will increase the presentation time.

To ensure the task was measuring surround suppression and ability to ignore movement of surrounding dots, a red circle appeared briefly before each trial indicating the location of the inner circle which moved randomly to one of six locations for each trial. For the peripheral surround condition a black circle of 6 degrees appears briefly and disappears when the moving dots appear, indicating the region of dots that should be ignored and the surrounding dots outside of that area is the concern for motion detection. The size parameters of the small and large stimuli were based on the research of Foss-Feig et al. (2013), Schauder et al., (2017) and Tadin et al., (2003) with the a small inner circle of 2° and a larger inner circle of 6° and all conditions consisting of an outer circle of 15°. It is important to note despite having the same parameters of these studies the design approach is different considering the present study is a 4AFC RDK whilst these studies used 2AFC drifting gratings.

### Procedure

Participants were invited to participate with an email providing a link to a qualtrics questionnaire along with debriefing and consent forms of this task. Once completed they were provided with the task file and a tutorial video on how to install Psychopy and perform the task. Upon installation of Psychopy, screen distance of 57cm must be maintained along with entering screen size and resolution to ensure presentation of stimuli was identical for each participant. A 60hz display must be used to ensure stimulus speed is not altered, a mac version was supplied to account for retina display. Once screen settings were configured the task commenced with instructions to use arrow keys to indicate the approximate direction that the majority of the dots are moving in. A 15 trial practice task provided feedback on correct and incorrect responses to confirm the participant understands the task. Each correct response decreased the amount of time the stimuli appeared allowing for a motion inspection time threshold to plateau within the 30 trials. Prior to each stimuli size change a screen prompted indicating a change in task, as well as for the peripheral surround condition.

The peripheral surround condition lasted for 60 trials in total and ended with a thank you participating screen to inform the participant the experiment has concluded. Psychopy generated the task data. Data was recorded automatically and generated within the data folder. The participants were instructed in the video to compress the folder to email to the researcher. Motion inspection time values were gathered from an excel sheet to plot onto a threshold graph.

**Figure 3.**
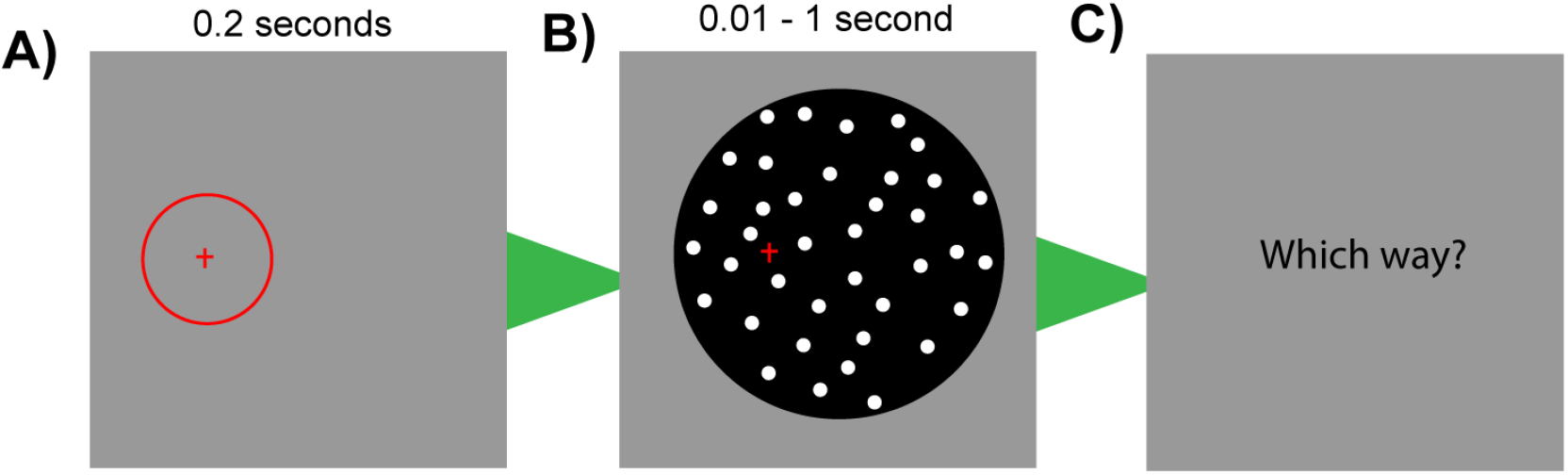
Schematic of Psychopy RDK motion task: **(A)** Participant must identify the location of the briefly appearing red circle. (**B)** Based on the location of the red circle, and with the assistance of the red cross visual cue, must observe the coherent direction (direction majority of the dots are moving in) the dots are moving in. The duration of this stimulus will depend on the amount of attempts guessed correctly (smaller duration for more correct guesses), ceiling effect will occur at 1 second and the minimum duration of 0.01 seconds. (**C**) When the RDK display disappeared, viewers were presented with text asking “Which way?, where they would have to respond with up, down or left or right keys.

### ANALYSIS OF PSYCHOPHYSICAL DATA

Inspection times were collected from the output excel documents, they were then labelled for each participant and their final inspection threshold was noted for each of the 6 conditions. Final inspection time was derived from 30^th^ trial for each condition. The AQ score provided from qualtrics was paired with their threshold times and a one way ANOVA was conducted on IBM SPSS 13.0 to find significance between autistic trait levels and motion inspection time. Descriptive statistics was conducted on SPSS on analyse population age, gender and Independent T test was used to test main effects between AQ score and inspection times. Scatter plot and Pearson’s correlation was conducted in SPSS to measure relationship between AQ scores and motion inspection times.

**Figure 4.**
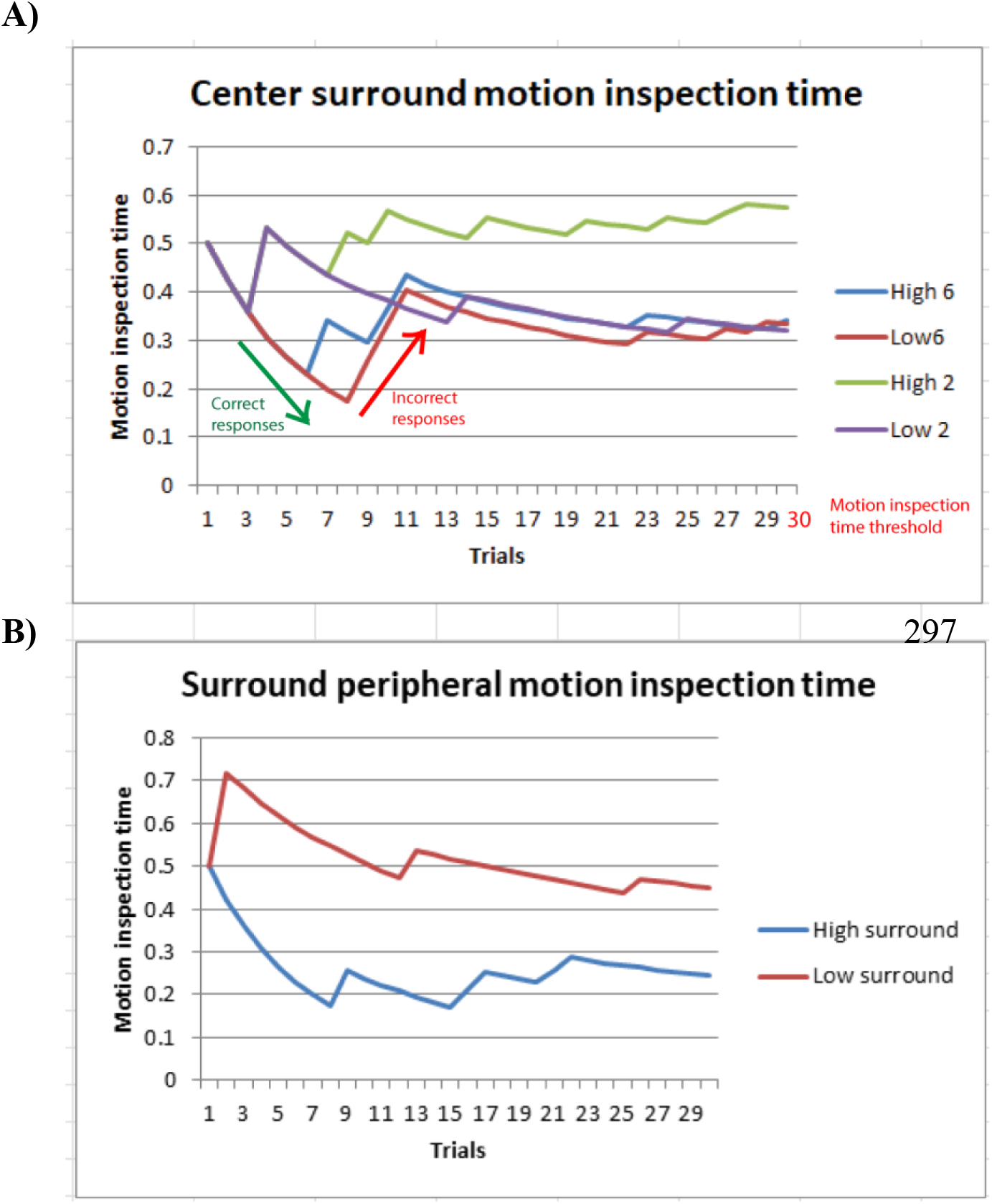
Individual’s attempt plotted to observe interleaved staircase QUEST and final inspection time threshold **A)** Demonstrates motion inspection time for conditions with small and large inner circles (center) at high and low contrast. Initial motion inspection time begins at 0.5 seconds at trial 1 with the final inspection time deduced from trial 30 **B)** Surround peripheral conditions, of high and 6° inner circle with 15 ° surround was used in both high and low contrast conditions.

## RESULTS

### Population assessment

Survey results indicated that of the 12 participants AQ scores ranged from 13 to 37 (M=21, SD=8.10). No participants presented with a low AQ score, 9 participants were labelled with a medium AQ score and 3 were labelled with a high AQ score. A binary split was conducted to split participants based on their AQ scores

**Table 1.** Participant demographic statistics

|  |  | Mid AQ (n=9) | High AQ (n=3) | Total (n=12) |
| --- | --- | --- | --- | --- |
| Gender | Male | 6 | 3 | 9 |
|  | Female | 3 | 0 | 3 |
| Age | 18 -25 | 9 | 2 | 10 |
|  | 26 - 35 | 0 | 1 | 1 |

**Table 2.** Motion inspection time for center coherence condition

|  |  | High contrast |  | Low contrast |  |
| --- | --- | --- | --- | --- | --- |
| AQ Group |  | 6 degrees | 2 degrees | 6 degrees | 2 degrees |
| Medium |  | 0.32 | 0.63 | 0.55 | 0.53 |
| High |  | 0.42 | 0.47 | 0.27 | 0.29 |

Note. Motion inspection time in seconds, lower inspection times indicate better performance

**Table 3.** Average Motion inspection time for surround coherence condition

| AQ Group | High surround | Low surround |
| --- | --- | --- |
| Medium | 0.59 | 0.42 |
| High | 0.44 | 0.44 |

*Note.* Motion inspection time in seconds

Between group analysis: A one-way analysis of variance (ANOVA) was conducted to assess high and medium AQ between size and contrast level of the RDK task (Table 4). In the centre high contrast condition no significance was found, along with the high and low contrast surround conditions. AQ score was found to be a predictor of the small low contrast condition (SD=0.26, p=0.04)

**Table 4.** One way ANOVA between AQ levels and motion inspection time threshold

| Condition | Contrast | Size | F | p |
| --- | --- | --- | --- | --- |
| Center | High | 6 Degrees | 2.70 | .49 |
|  |  | 2 Degrees | 4.51 | .53 |
|  | Low | 6 Degrees | 5.32 | .019* |
|  |  | 2 Degrees | 5.68 | .030* |
| Surround | High | 6 Degrees | .24 | .93 |
|  | Low | 6 Degrees | .63 | .76 |

*Note.* \* $p < 0.05$ = Significant

**Figure 5.**
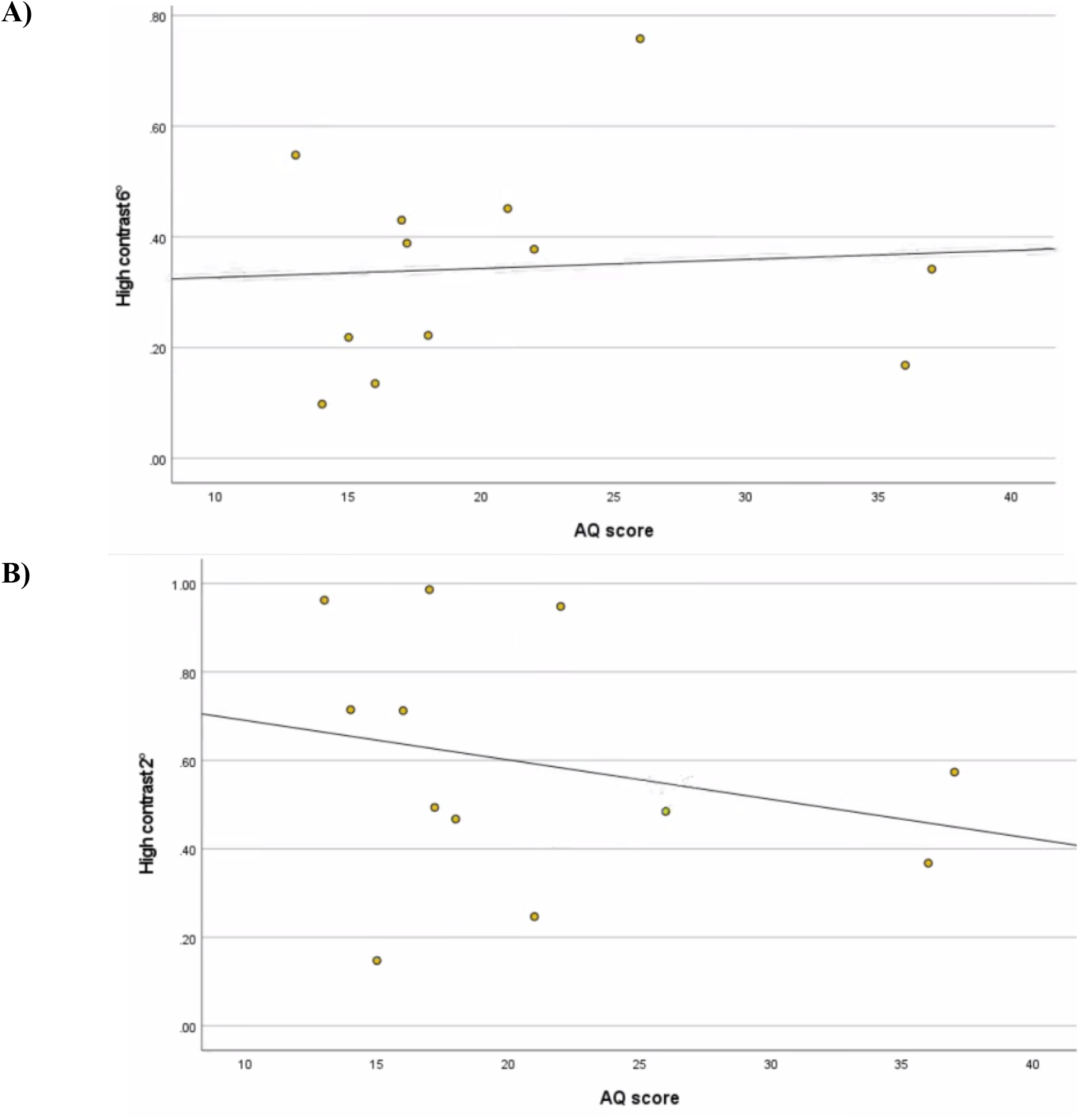
Scatter plot of large (**A**) and small (**B**) high contrast stimuli motion inspection time for 12 participants based on their AQ score. Mid AQ group in this sample are scored 13-22 (n=9), high AQ group are scored 26-37 (n=3)

A Pearson correlation was conducted for small and large high contrast conditions and AQ score and found a very weak positive relationship was found for large stimuli size (r=.07,p=.83) and not statistically significant. For the small sized stimuli a weak negative relationship was found (r=-.26, p=.41) and not statistical significance. The high AQ group motion inspection time was on average higher (M=0.76, SD= 0.30 seconds) than the mid AQ group (M=0.55, SD=0.15 seconds) for the large high contrast condition with a 6 degree inner circle, indicating the mid AQ group performed better requiring less time to view the stimulus. For the small high contrast condition high AQ group (M=0.47, SD=0.10 seconds) outperformed the mid AQ group (M=0.63, SD= 0.31 seconds). However, Independent sample T test was conducted between high and mid AQ group for both large (t=0.71, p=0.49) and small (t=0.66, p=0.53) high contrast and no significance was found.

**Figure 6.**
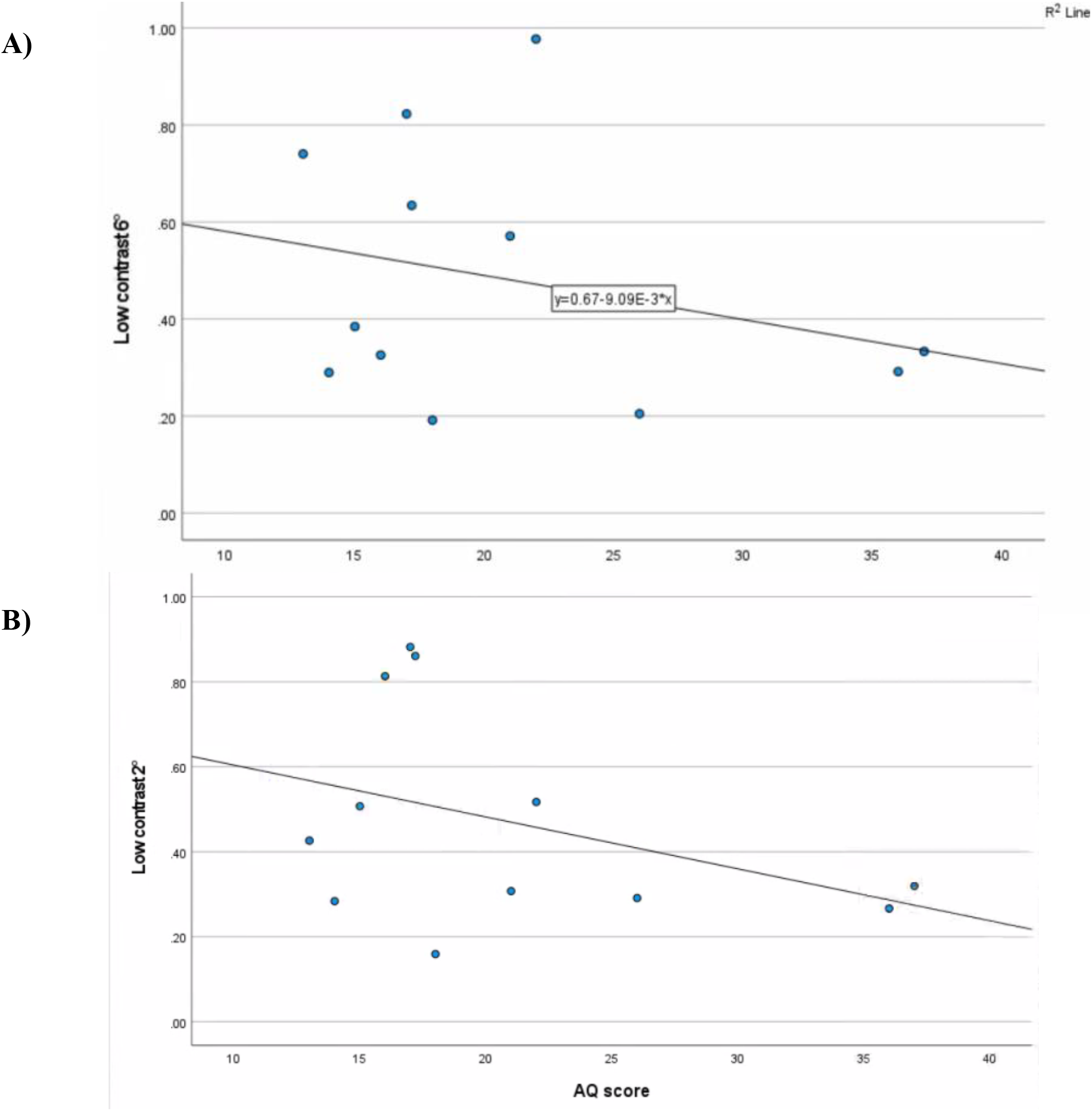
Scatter plot of large **(A)** and small (**B**) low contrast stimuli motion inspection time for 12 participants based on their AQ score

A Pearson correlation was conducted for small and large low contrast conditions and AQ score and found a weak negative relationship was found for large stimuli size (r=-.28,p=.38) and not statistically significant. For the small sized stimuli a weak negative relationship was found (r=-.39, p=.21) and no statistical significance.

An Independent t test was conducted and found that low contrast condition showed significance differences between high and mid AQ group motion inspection times for both small (t=2.43, p=0.36) and large (t=2.80, p=0.19) stimuli. The high AQ group (M=0.27, SD=0.66 seconds) out performed mid AQ group (M=0.55, SD= 0.26 seconds) for large condition and for the small condition the high AQ group (M=0.29, SD= 0.26) also outperformed the mid AQ group (M=0. 55, SD= 0.26 seconds).

**Figure 7.**
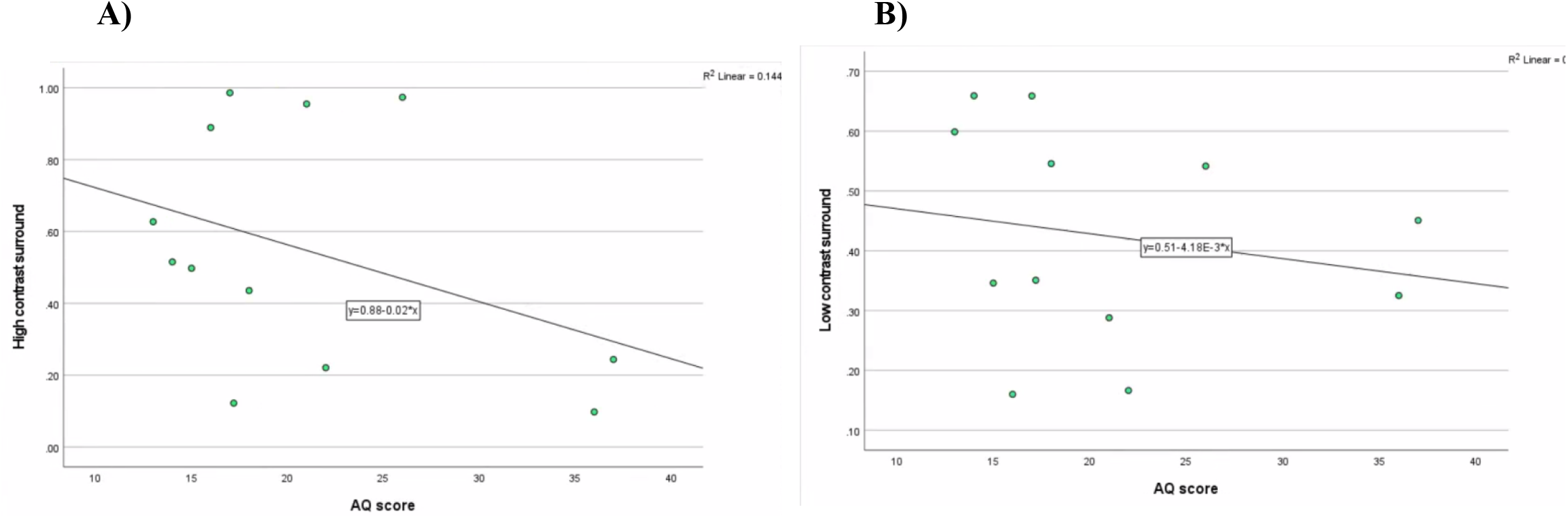
Scatterplot of high (**A**) and low (**B**) contrast surround motion inspection time for 12 participants based on their AQ score. Mid AQ group in this sample are scored 13-22 (n=9), high AQ group are scored 26-37 (n=3)

A Pearson correlation was conducted for high and low contrast surround conditions and AQ score and found a weak negative relationship was found for high contrast (r=-.38,p=.22) and not statistically significant. For low contrast a moderate positive relationship was found (r=.55, p=.22) and not statistically significant.

On average the high AQ group (M= 0.44. SD= 0.47) outperformed the mid AQ group (M=0.58, SD= 0.31 seconds) for the high contrast surround condition. The mid AQ group (0.42, SD= 0.20 seconds) outperformed the high AQ group (M=0.44, SD=0.11 seconds).

## DISCUSSION

The study aimed to investigate whether level of autistic traits affects performance of motion perception tasks involving varying size and contrast. The results are inconsistent with previous studies, irrespective of condition, no deficits in gain control for the high AQ group, and overall improved performance across all conditions for high AQ was observed. However, high AQ was only found to be a predictor of performance in small and large low contrast conditions. Stimuli size did not show significant delays in motion inspection time for either mid or high AQ groups, implicating that surround suppression did not alter performance of typically developed individuals.

The variance in our results opposing Tadin’s work could be accounted by the relationship between spatial and temporal factors in surround suppression (Churan, Richard & Pack 2009; Alitto & Usrey 2015). Churan, Richard & Pack (2009) adds the functionality of receptive fields residing within the LGN are influenced by duration of stimuli along with the spatial structure of stimuli, in turn mitigating the mixed results often seen in psychophysical studies concerned with centre surround suppression.

The RDK design used in this study presented a brief static visual cue for 2 milliseconds prior to the motion stimulus to assist with orienting attention and reduce search time. This is an imperative element of the task as the stimulus switched between to 6 random locations. This switching discourages focusing on a central visual cue and assess how surround suppression occurs with a subtle form of spatial uncertainty, however the static target used prior to dot appearance may remove the counterintuitive effects of stimulus size (Churan, Richard & Pack 2009). Previous study state this occurs due to the problem of receptive fields tuning to orientation of a static stimulus and readjusting to a moving stimulus, discounting the effect of size on perception of briefly presented motion displays (Borghuis et al., 2019).

Other parameters to take into account that were manipulated throughout this study were dot life time, dot size and speed. Manning, Charman & Pellicano (2015) mentioned that ASD individuals performed better in dot life times that were unlimited, dot life time remained constant at an unlimited level for all conditions throughout the study. However instead of measuring motion inspection time limited dot lifetime may be used as a means of task difficulty, along with motion coherence levels to assess local vs global multiple object tracking (MOT) as opposed to motion inspection time (Manning, Charman & Pellicano 2015; Koldewyn et al. 2012).

The parameter motion coherence level was set to 50% in this study allowing for visual noise. Studies have shown ASD invididuals are influenced by the presence of noise however still demonstrate intact multi-sensory (visual-vestibular) integration (Zaidel, Goin-Kochel & Angelaki 2015). This can account for unaffected eye movements seen in ASD individuals regardless of delayed or enhanced performance in motion inspection times. Furthermore some participants reported being unable to detect motion coherence at 50%, suggesting either 0.5 seconds does not allow them enough time to distinguish motion, the visual of cue of 0.2 seconds did not register in their endogenous attention to search for motion within the RDK, or the combination with visual noise and stimuli size makes the center and surround dots undistinguishable.

Modulation of attention is likely to be a key factor interacting with results, taking into account the target stimuli switched randomly between 6 locations for each trial eliciting spatial uncertainty. Furthermore, the ability to sustain attention was problematic for a few of the participants that achieved ceiling effect. Spatial uncertainty is a variable that has been explored in previous work, with results suggesting that larger attentional fields emerges for stimuli with spatial uncertainty. This was not measured in the present study, implicating measure of response gain had possibly masked the effects of gain control and surround suppression (Herrmann et al. 2010). The ceiling effect seen among 4 participants are suspected to have such results due to this phenomenom. Their motion threshold exceeded the initial stimulus presentation of 0.5 seconds meaning they had to be excluded as the true motion inspection times of surround suppression and gain control did not factor into their performance. Endogenous attention (understanding of task) must be taken into account to allow for exogenous attention (driven instinctively) especially considering a fast paced task involving fast adaption of motion perception as studies have shown slow exodenous orienting in ASD.

Past studies have also speculated that abnormal attentional deployment may relate to receptive fields sizes along with impaired response gain, requiring more time to fixate on a moving stimulus (Schauder et al. 2017; Sprague & Serences, 2013). In addition to attention deployment, other confounds in the results could be explained by eye movements and binocular rivalry. (Arranz-Paraíso et al., 2021; Robertson & Baron-Cohen., 2017).

It is commonly discussed within the ASD literature that ASD individuals struggle to extract social information from biological motion, in addition presenting with behavioural tendencies presenting in the form of reduced eye contact and or abnormal eye gazes while processing complex stimuli (Robertson & Baron-Cohen., 2017; Madipakkam et al., 2017). Furthermore saccade eye movements (rapid movement between points) have been shown to modulate receptive field responses (Takarae et al., 2013). This higher order processing occurs by spatial sensitive neurons in the V4 serving a crucial role in discerning and scanning for points of interest in the environment (Tolias et al., 2001; (Takarae et al., 2013).

Robertson & Baron-Cohen (2017) discuss ASD individuals often demonstrate faster detection of single details surrounded by visual distractions. Furthermore, ASD deficits are visible specifically when viewing global visual displays, the motion signal is weak and the amount of time to integrate the movement of stimuli is short. In conjunction with variance in ASD motion processing that previous studies have noted, clinical populations were previously used in their studies, in contrast this study grouped consisted of mostly typically developed individuals. The highest AQ score (26) not including the one individual diagnosed with autism (AQ of 36) do not appear to show any form of deficits throughout all conditions, rather improved performance especially in low contrast conditions. Observing the performance of the diagnosed ASD individual, such high performance could implicate hypersensitivity in their visual system along with regular exposure video games compared to the rest of the sample.

A number of limitations were encountered during this study, requiring attention for future studies. It is important to consider the questions that provided the ASQ scores derived from the questionnaire were purely self-assessed, allowing participants for untruthful/incorrect input. Furthermore binary grouping was used in determining levels of autistic traits that cannot necessarily be generalised for clinical population. Taking into account the study only recruited 12 participants, 3 being high AQ and 9 being mid AQ, making it difficult to assume whether performance is based on individual differences or neurological/sensory functioning related to autistic traits. Larger samples will allow for comparison of low AQ vs high AQ scorers and finding significant trends on whether the size and contrast influences inspection time rather than unknown confounding variables within a small sample influencing the results. Matched cohort design with younger children will be crucial for future work studying motion perception, considering maturation of visual pathways have not fully matured compared to older age groups (Shuffrey et., 2018) Understanding of the task may have played part as the experiment was delivered online, without the supervision of a researcher or assistance to perform the task. Visual acuity tests and assessments involving detection of stimuli with no noise (100% motion coherence) should also be assessed prior testing to evaluate whether the individual is able to detect or see moving stimuli with the default parameter settings, to ensure visual deficits unrelated to ASD are not impacting performance.

The stimuli presented were not consistent due to constraints of using Psychopy 3.0. Dot density parameter was limited to whole numbers rather than decimals resulting in either underpopulated or overpopulated moving dots for the small and large conditions. This was clearly visible in the practice trials where the participant were met with dots that were inconsistently spaced and difficult to perceive in small conditions but very easy to perceive in large conditions, causing disparity in motion inspection time thresholds. For this reason 30 trials per condition were established rather than randomly mixing the conditions. The study did not utilise eye tracking like previous studies did, doing so may show underlying processes in regards to hierarchical attention, retinal slip and gaze patterns. Eye tracking may not be prevalent in typically developed individuals for the RDK paradigm used in this study, however in a large clinical sample of ASD participants; this is a factor to take under consideration. Falck-Ytter et al. (2013) emphasises that eye tracking in ASD can provide beneficial data with comparison of spatial and temporal gaze patterns. Furthermore the study design could be altered, rather then having 30 trials of each condition (contrast and size) mix all the conditions together and observe how higher autistic trait group approach the switching of conditions (Dakin and Frith, 2005). Eye tracking may measure adaptability and hierarchical approach to spontaneous switching of conditions, whether there is a cognitive bias to focus more on certain stimuli contrasts and sizes. In addition to eye movements, surround suppression seems to occur primarily in binocular (both eyes) viewing of stimuli, more so than monocular viewing. The notion that MT cells are binocular raises questions whether the high prevalence of strabismus in ASD (crossed eyes) may contribute to reduced surround suppression (Arranz-Paraíso, Read & Serrano-Pedraza 2021).

The current study provided new insight into possible masking effects of stimuli size and enhanced performance of low contrast irrespective of stimuli size. To our knowledge this is the first study to conduct a 4 AFC RDK over a 2 AFC with both center and surround conditions of varying size and contrast to assess surround suppression grouped by level of autistic traits. The implementation of a visual cue prior to motion displays, allowed us to remove of stimuli borders that would have confounded the ability to measure surround suppression. In summary the experimental stimuli design used did not present with delayed times for lower autistic traits in large high contrast conditions, suggesting the improved performance seen for high AQ scorers in low contrast does not demonstrate impaired gain control or surround suppression. Rather spatial and temporal interaction motion inspection time performance in low visibility conditions. Given the higher prevalence of sensory deficits in ASD, understanding the vulnerabilities the dorsal pathway poses on visual and motion perception is crucial to underpinning the behavioural tendencies that ASD individuals display.

